# Forest fire survival in young, dense *Betula ermanii* stands on soil scarification sites

**DOI:** 10.1101/2020.09.20.305557

**Authors:** Masato Hayamizu, Yasutaka Nakata, Hiroyuki Torita

## Abstract

A forest fire in a cool-temperate broad-leaved forest in northern Japan, from 26 May to 19 June 2019, provided an opportunity to examine its effects on young and dense birch (*Betula ermanii* Cham.) stands in soil scarification sites. To characterise post-fire responses (survival and resprouting) of birch, we set up two plots, 6 months post fire. We investigated trunk diameter at breast height and burn marks on tree trunks (scorch height and charring percentage around the tree bole) of all *B. ermanii* trees in both plots. Survival and resprouting of each tree were monitored over 2 years (6 and 16 months post fire). To quantify post-fire vegetation recovery in the forest floor, we manually mapped the dominant understory plant, dwarf bamboo (i.e. *Sasa kurilensis* (Rupr.) Makino et Shibata), from orthomosaic images obtained by an unmanned aircraft vehicle, and estimated the recovery rate in the second year post fire. Additionally, seedlings of woody species were counted in both plots. Size-dependent survival rates of plants in both plots were similar in the first year post fire. All *B. ermanii* trees died without resprouting in the second year post fire, indicating the lethal effects of fire on young birch trees. Moreover, a high recovery rate of dwarf bamboos over 2 years in both plots and limited seedling establishment of woody plants suggest that the fire resulted in regeneration failure of young stands in the scarification sites. On the basis of these findings, we propose future management of stands in soil scarification sites post fire, considering the vulnerability of young trees and the rapid change in vegetation from young forest to dense birch cover post fire.

## Introduction

Fire is a common natural disturbance in forests worldwide and affects the dynamics of vegetation in various forest ecosystems (Bowman *et al.*, 2009). In fire-prone ecosystems, post-fire natural regeneration enables the persistence of forest vegetation, even after the destruction of large number of trees (Fernandes *et al.*, 2008; Catry *et al.*, 2013). However, the survival capacity of plants is not well understood in less fire-prone ecosystems, owing to limited opportunities to examine the process of post-fire vegetation dynamics (Goto *et al.*, 1996). Moreover, during recent decades, forest fires have been increasing due to human activities and climate change (Flannigan *et al.*, 2013; Keeley *et al.*, 2019). Therefore, understanding post-fire vegetation dynamics in less fire-prone ecosystems is as important as it is in fire-prone ecosystems (Tepley *et al.*, 2018).

Many plant species typically regenerate via two adaptive strategies post fire: seedling generation from seeds (Keeley and Fotheringham, 2000) and resprouting from roots and stumps (Bond and Midgley, 2001; Pausas and Keeley, 2014). Previous studies in fire-prone forests have demonstrated rapid recovery post fire through a flush of germination (Roy and Sonie, 1992; Tyler, 1995) and rapid vegetative resprouting (Catry *et al.*, 2013; Stevens-Rumann and Morgan, 2016). In cool-temperate broad-leaved forests in northern Japan, where plants are not adapted to fire (Nakagoshi *et al.*, 1987), forest vegetation can still regenerate post fire (Masaka *et al.*, 2000; Goto, 2004). In particular, post-fire seed production and resprouting involves large and mature trees (Masaka *et al.*, 2000). Higher canopies and thicker barks make larger trees more resistant to the heat of fire than smaller trees, and thus, larger trees are more likely to survive (Gill and Ashton, 1968; Pausas, 2015). This resilience of larger trees to heat damage is manifested as their ability to allocate resources within seeds, enabling resprouting post fire (Masaka *et al.*, 2004). On the contrary, young forests with small trees are considered more vulnerable to fire and less resilient than mature forests. However, there are only a few studies on the response of cool-temperate broad-leaved forests to fire disturbances, especially those focusing on post-fire survival and regeneration in young forest stands.

Another cause for the failure of natural regeneration is the inhibition of seedling establishment and growth through dense understory plants (Watt, 1919; Mallik, 2003). Several plant species can rapidly recover post fire, cover the forest floor faster than that under forest establishment, and demonstrate sprouting of tree seedlings in fire-prone coniferous forests (Mallik, 2003). In cool-temperate broad-leaved forests, the ground cover is frequently dominated by a dense evergreen understory of dwarf bamboos, *Sasa kurilensis* (Rupr.) Makino et Shibata (Masaki *et al.*, 1999, Matsuo et al. 2018). In northern Japan, dwarf bamboos reproduce vegetatively from their root systems to form a dense cover after less severe disturbances (Goto, 2004), which inhibit seedling emergence and survival of tree species (Taylor and Zisheng, 1992; Noguchi and Yoshida, 2004). Since the 1970s, soil scarification, by which both understory dwarf bamboos and surface soil are removed using engineering machinery, has been widely applied for assisted natural regeneration in natural forest management in northern Japan (Umeki, 2003; Ito *et al.*, 2018). More than 45,000 hectares of soil scarification sites have been established in Hokkaido, northern Japan (Ito *et al.*, 2018). Moreover, Umeki (2003) reported that 122 (84%) of the 146 stands surveyed in scarification sites in Hokkaido were young forest stands (13–22 years). Therefore, most of the stands in soil scarification sites in Hokkaido are still expected to be young forests, less than 30 years old.

*Betula ermanii* Cham. is a major broad-leaved tree species in cool-temperate mixed forests in northern Japan (Kikuzawa, 1988). *B. ermanii* is considered a pioneer species with disturbance-related characteristics such as high dispersal ability and fast growth potential in disturbed sites (Kikuzawa, 1988). It is the most dominant species and forms monospecific, even-aged stands in soil scarification sites (Umeki, 2003). Moreover, this species has the ability to sprout from the base of the trunk and is known to play an important role in the persistence of individuals under severe environments (Okitsu, 1991). However, little is known about the responses of young birch trees post fire disturbance (especially on survival and germination) and the responses of forest floor vegetation that affects post-fire natural regeneration of birch trees.

In Hokkaido, large-scale fire disturbances occurred at a higher frequency before the start of the 20^th^ century (Takaoka and Sasa, 1996), but according to the Japan Ministry of Agriculture, Forestry and Fisheries, relatively small-scale fires continue to occur sporadically. Owing to the humid temperate regions in northern Japan, the intensity of fires is estimated to be within the range of surface fire line intensities, as reported in forests in the United States and Canada (Goto *et al.*, 2005). Although most of the young forest stands in soil scarification sites in Hokkaido have not yet experienced fires, we hypothesised that the vulnerability of young trees to catch fire and the resilience of the dwarf bamboos result in regeneration failure of cool-temperate broad-leaved tree species.

A forest fire that started in Ōmu Town, north-western Hokkaido in May 2019 burned 214.79 ha of forest. This was the largest surface fire in Hokkaido in terms of area and scale in the past 30 years (Hokkaido Government, 2019). This provided an opportunity to examine the effects of fire on the survival of young birch forests and examine the recovery of understory vegetation dominated by dwarf bamboo. In this study, post-fire responses of *B. ermanii* trees in scarification sites were evaluated to examine the survival and resprouting of dense, young birch forests. The objectives of this study were to: (1) characterise the post-fire responses, especially survival and resprouting, of *B. ermanii* trees and (2) quantify vegetation recovery in the forest floor post fire. Here, we discuss the post-fire natural regeneration of *B. ermanii* stands in soil scarification sites.

## Materials and Methods

### Study sites

The study sites are located in the scarification sites in Ōmu Town (Figure 1a and 1b). A fire was first detected on 26 May 2019 and was finally declared to be under control in this area on 19 June 2019. The vegetation on the forest floor in the study areas was almost completely destroyed by the fire (Figure 1c and 1d). The fire spread across 214.79 ha, comprising 165.83 ha of a secondary cool-temperate broad-leaved forest and 48.96 ha of a plantation coniferous forest. There are no records of previous fires in these study sites (Hokkaido Government, 2019). The weather during the fire was sunny, with a mean temperature of 25.5°C (maximum temperature; 32.4°C), mean humidity of 39%, maximum wind velocity of 13.8 ms^-1^, and maximum instantaneous wind speed of 24.2 ms^-1^, with a southeast wind direction (Japan Meteorological Agency, 2019). Ōmu Town belongs to the Okhotsk climatic zone, and the forest floor is usually snow-covered from December to April in the subsequent year. In May, the frequency of high-temperature events was higher in Okhotsk area relative to that in other areas (Mori and Sato, 2014). Combined with droughts caused by the foehn wind, the surface fire fanned by the foehn winds burned the study area soon after the disappearance of snow cover (Mori and Sato, 2014; Hokkaido Government, 2019). The elevation of the site ranges from 500 to 560 m. Soil scarification was conducted in 1989, and the age of the *B. ermanii* forest stands was 30 years or less in 2019.

**Figure 1.**
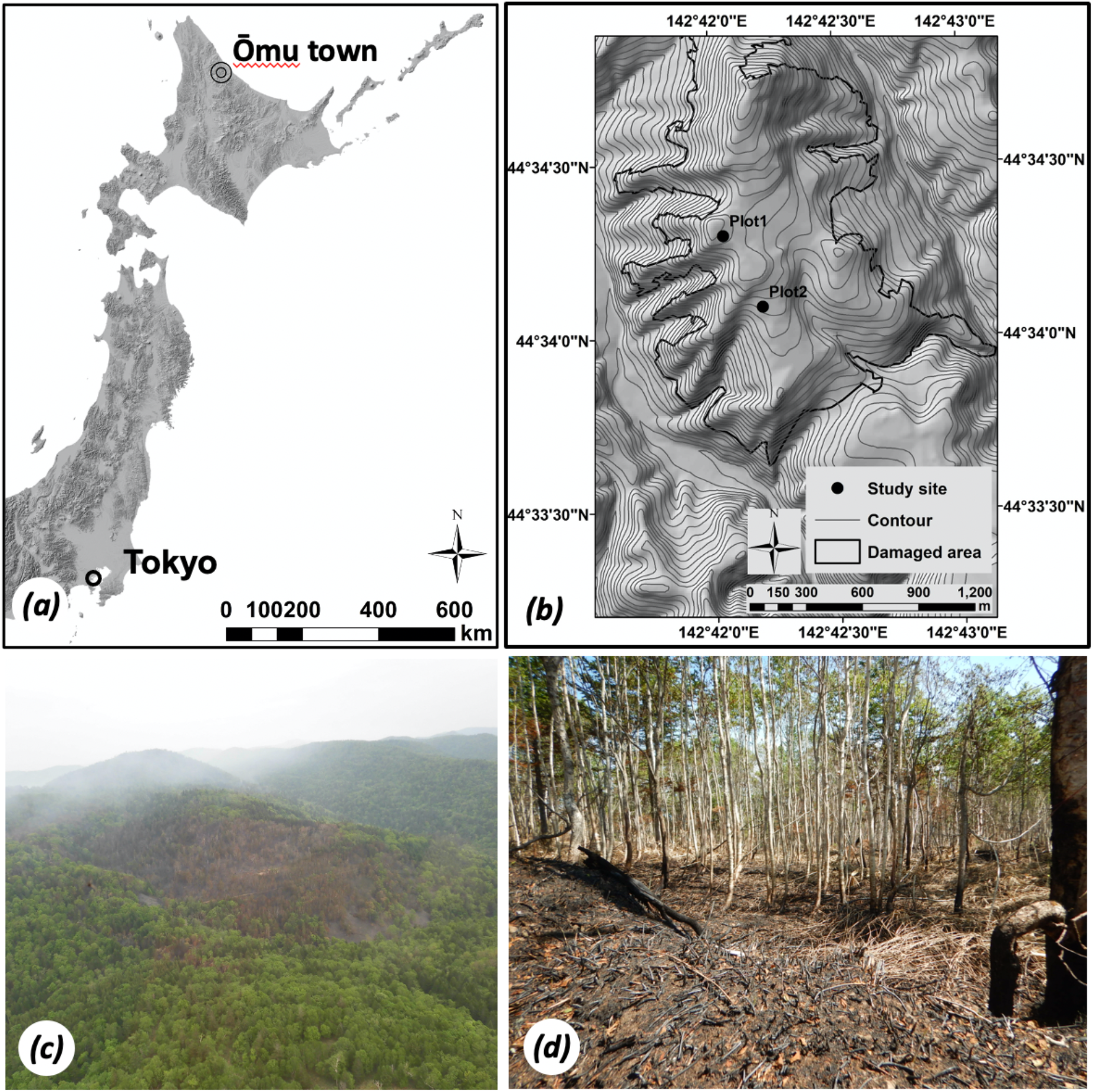
Map and location of Ōmu Town, Hokkaido, Japan. Location of the two study sites (a). Black dots represent the study plots, and the area surrounded by the black line indicates the burnt area traced during a field survey (b). Aerial view of the scene immediately after the fire (c). *Betula ermanii* forest stands near the plot immediately after the fire (d).

### Field survey and data collection

In the forest fire sites, two plots of burned *B. ermanii* stands were established in scarification sites on 30 October 2019. In Plot 1, which was scarified in a rectangular shape in 1989, a 10 m × 50 m quadrat was set up on a gentle southeast-facing slope at an altitude of 560 m. In Plot 2, which was scarified in a square shape in 1989, a 20 m × 20 m quadrat was set up on a gentle south-facing slope at an altitude of 503 m (Figure 2).

**Figure 2.**
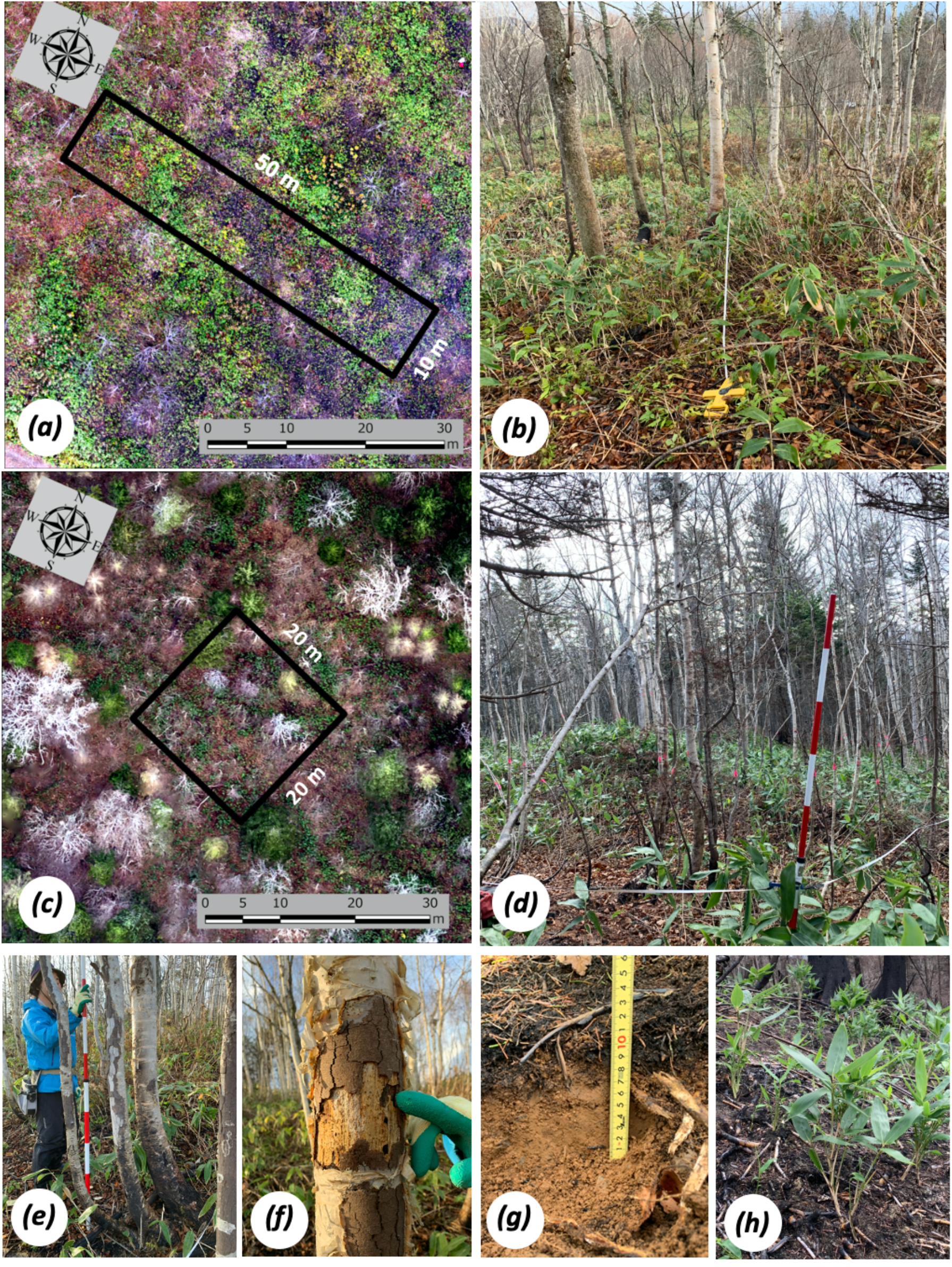
*Betula ermanii* stands in scarification sites after 6 months in Plot 1 (a-b) and Plot 2 (c-d). Each plot was set to the position indicated by black squares (a, c). Scorched tree (e); burnt and damaged tree (f); litter layer and burnt litter (g; black layer); and understory vegetation and recovered *Sasa kurilensis* (the most dominant species) (h).

To examine tree mortality and resprouting after a fire, only trees with a diameter at breast height (DBH) of above 0.7 cm were sampled; overall 112 *B. ermanii* trees were measured and observed in Plot 1 and 115 birch trees were measured and observed in Plot 2. Data were collected from 30 October to 1 November 2019 (6 months post-fire) and on 10 September 2020 (16 months post-fire). Trees in the study plot were tagged, their DBH was measured to the nearest 1 cm at 1.3 m above the ground level, and new sprouts were counted. Tree size measurements included the DBH for all birch trees (Plot 1, *N* = 112; Plot 2, *N* = 114; Table 1). Tree height and clear length of randomly selected representative trees in the plot were measured (Plot 1, *N* = 6; Plot 2, *N* = 10; Table 1). Tree survival was divided into binary categories by visual inspection of crown condition: 0, dead, no foliage or sprouts in the crown; 1, alive, foliage survived, and sprouts or inflorescence buds were present in the crown. Scorch height and bole charring percentage (stem damage), which is correlated with fire intensity, of all individuals were measure. The values for the plot and transect variables (Table 1) were used to characterise all trees in a given plot.

**Table 1:** Summary of the structural properties and assessed variables of *Betula ermanii* stands in scarification sites

|  | Area | density | DBH | Tree height | Clear length | Litter depth | Scorch height | Stem damage | Burned<br>litter depth |
| --- | --- | --- | --- | --- | --- | --- | --- | --- | --- |
|  | (m) | (ha<sup>-1</sup>) | (cm ± SE) | (m ± SE) | (m ± SE) | (cm ± SE) | (m ± SE) | (% ± SE) | (cm ± SE) |
| Plot 1 | 10 × 50 | 2240 | 5.32 ± 0.35 | 7.77 ± 0.73 | 3.64 ± 0.33 | 2.79 ± 0.77 | 1.49 ± 0.10 | 62.79 ± 2.88 | 1.39 ± 0.38 |
| Plot 2 | 20 × 20 | 2875 | 5.51 ± 0.28 | 7.71 ± 0.71 | 4.08 ± 0.32 | 4.27 ± 1.16 | 0.68 ± 0.03 | 87.02 ± 1.45 | 0.96 ± 0.19 |

DBH: Diameter at breast height, SE: Standard error

To estimate the extent to which the heat of the fire reached the belowground parts of the plant, burned litter depth was recorded in 1 m × 1 m quadrats, randomly repeated three times in each plot, and was measured as the depth to which the burn marks reached in the litter layer, after measuring the depth of the litter layer (litter depth) of the plot. All canopy tree was recorded in each plot, 16 months post fire. Understory plants, especially seedling and other herbaceous plants, were recorded in 1 m × 1 m quadrats, randomly repeated three times in each plot, 16 months post fire.

### Monitoring of vegetation recovery

To estimate the recovery of dwarf bamboos, an unmanned aircraft vehicle (UAV; real time kinematic-UAV (RTK-UAV) Phantom 4 RTK (DJI Co., Shenzhen, China)) was used. The device was equipped with a D-RTK 2 high precision global navigation satellite system (GNSS) mobile station, which allowed real-time differentiation between the aircraft and the local GNSS system, or an RTK base station. The flight height was set to 100 m to acquire high-resolution orthorectified images. These images were created using the structure-from-motion and multi-view stereo algorithms. The algorithms automatically detect feature points to match with two-dimensional digital images. Based on the detected feature points of the target, three-dimensional spatial information of the target was acquired, and then an orthorectified image was created with this information. Metashape version 1.5.3 (Agisoft LLC, Saint Petersburg, Russia) was used to process the photographs. Ground truth and vegetation properties were recorded for all plant species by name. Culm height of the dwarf bamboos was directly calculated in plots 1 and 2, 16 months post fire.

### Data analysis

The data were analysed using generalised linear models (GLMs). All statistical analyses were carried out using the statistical software R, version 3.6.2 (R Development Core Team, 2019). As dependent variables, we used post-fire tree responses, especially individual mortality, that is, mortality of all aboveground and belowground organs (tree death); these post-fire responses were examined in relation to different explanatory variables collected at the individual level (Table 1). For each response variable assessed, we started with a model including all the explanatory variables. Selection for the GLMs was performed with the R package ‘MuMIn’ to select the minimum Akaike information criterion (AIC) value, with differences in AICc (ΔAICc) >2 used as evidence of substantial model dissimilarity (Barton, 2013).

In both plots, the cover and recovery rates of the dwarf bamboos were calculated by manual mapping using RTK-UAV orthomosaic images at two time points, 6 and 16 months post fire. The UAV orthoimages were mapped and distinguished manually by a human interpreter in the GIS using ArcGIS software version 10.6 (ESRI Inc., Redlands, CA, USA).

## Results

### Tree survival and resprouting

The 227 surveyed trees showed a clear size-dependent high mortality rate (Table 2, Figures 3 and 4). Post-fire mortality was 75.9% in Plot 1 and 72.2% in Plot 2 at 6 months post fire. Despite careful observation, post-fire mortality was 100% in both plots at 16 months post fire. In Plot 1, with a modal DBH of 3 cm and stand density of 2240 trees ha^-1^, all plants with a DBH less than 5 cm died. The mean DBH of dead individuals was 5.32 ± 0.35 cm (mean ± standard error (SE)). The DBH of surviving individuals showed a modal value of 10–12 cm with a mean DBH of 10.57 ± 0.69 (mean ± SE). The survival rate was 24.1%, and the mean DBH of the surviving trees was significantly higher than that of the dead trees (*t*-test, *P* < 0.01; Table 2). The survival rate of trees in Plot 2, with a stand density of 2875 trees ha^-1^, showed the same tendency as in Plot 1. The modal DBH was 3 cm, and all individuals with a DBH of less than 5 cm died. As in Plot 1, resprouting from the burnt bases of *B. ermanii* stems was not observed in Plot 2. The mean DBH of the dead individuals was 4.18 ± 0.20 cm. The DBH of the surviving individuals was mostly 7–8 cm, with a mean DBH of 8.97 ± 0.42 cm. The survival rate was 27.8%, and the mean DBH of the surviving individuals was significantly higher than that of the dead trees (*t*-test, *P* < 0.01; Table 2).

**Figure 3.**
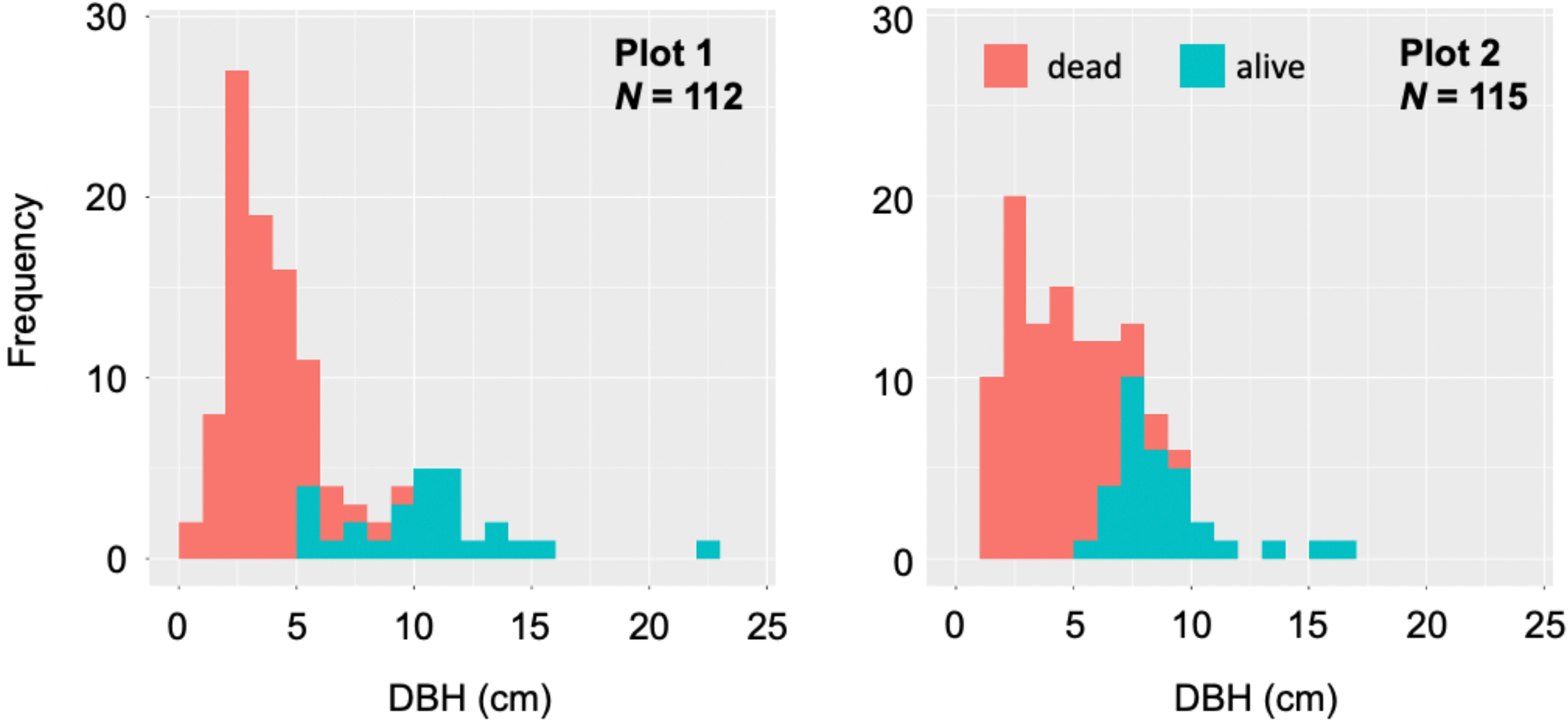
Stand structure of *Betula ermanii* in Plots 1 and 2. Histograms were created from individual trunk diameter at breast height (DBH) for each plot.

**Figure 4.**
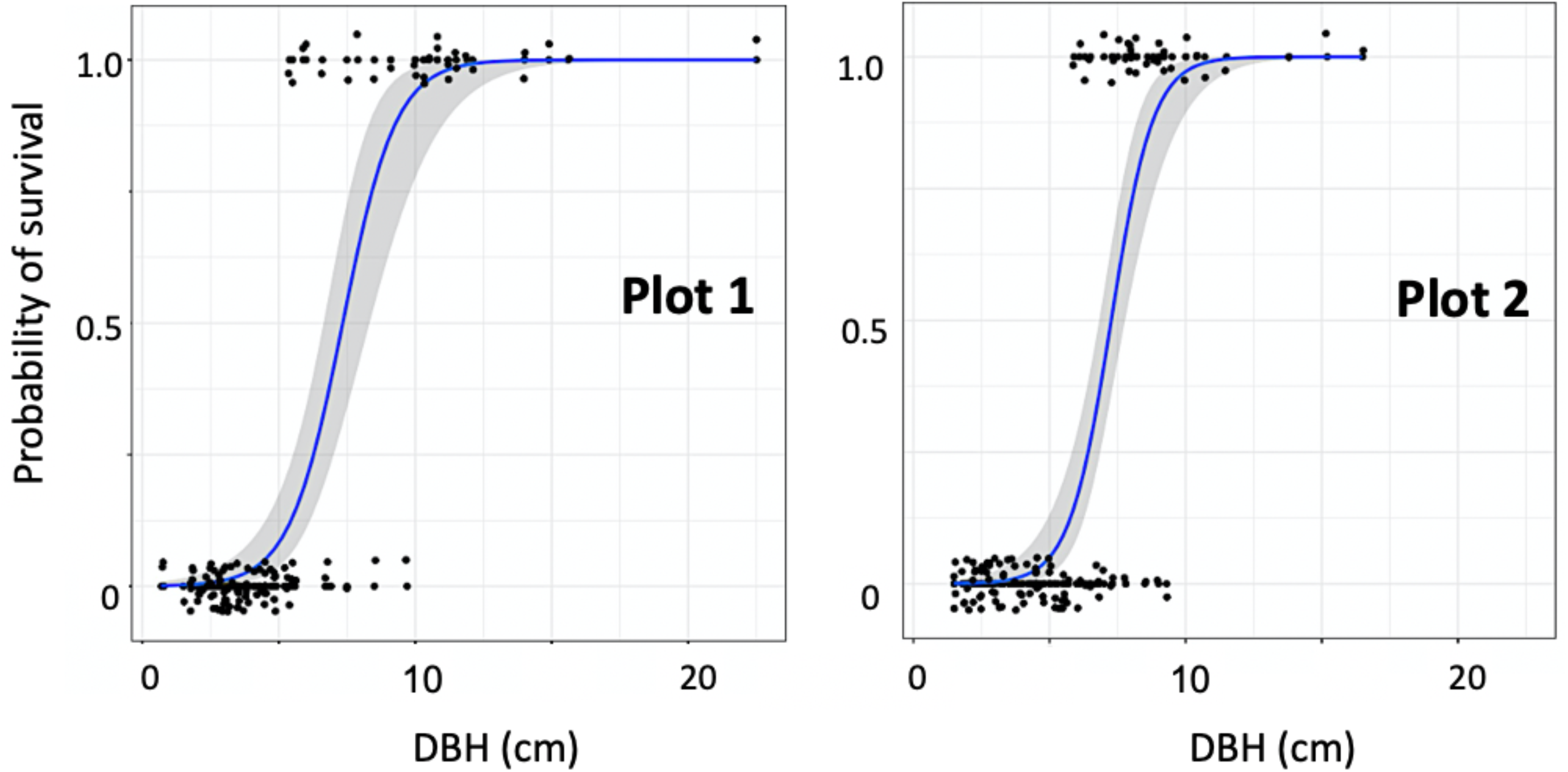
Binary logistic regression analysis of *Betula ermanii* as a function of trunk diameter at breast height (DBH). Black points around the ‘1’ line on the y-axis indicate surviving trees and black points around the ‘0’ line indicate dead trees. The solid blue line represents the probability function derived from the prediction equation and the grey area shows the 90% confidence interval.

**Table 2:** Summary of individual *Betula ermanii* survival at 6 months post fire.

| | | *N* | DBH mean ± SE (cm) | Survival rate (%) |
| --- | --- | --- | --- | --- |
| Plot 1 | Alive | 27 | 10.57 ± 0.69 <sup>a</sup> | 24.1 |
| | Dead | 85 | 3.65 ± 0.17 <sup>b</sup> | |
| Plot 2 | Alive | 32 | 8.97 ± 0.42 <sup>a</sup> | 27.8 |
| | Dead | 83 | 4.18 ± 0.20 <sup>b</sup> | |

4 Means within a column followed by different superscript letters (a and b) are significantly 5 different ( $P < 0.01$ ) for live and dead birch trees, respectively, as determined using a *t*-test.
DBH: diameter at breast height, SE: standard error, *N*: absolute number of individuals

In the GLM analysis, DBH showed significant effect on probability of survival of *B. ermanii*, whereas scorch height and stem damage were not significantly different from one another (Table 3). The probability of survival of *B. ermanii* showed significant dependence on DBH in both plots (Figure 4), which was consistently included in the best models (Supplemental Table 1).

**Table 3:** Generalized linear model (GLM) analyses of the effect of probability of survival of *Betula ermanii*.

|  | Estimate | SE | z value | *P* value |
| --- | --- | --- | --- | --- |
| Intercept | -9.142 | 1.731 | -5.283 | <0.001* |
| DBH | 1.158 | 0.173 | 6.678 | <0.001* |
| Scorch height | 0.001 | 0.003 | 0.291 | 0.771 |
| Stem damage | 0.008 | 0.012 | 0.666 | 0.505 |

Asterisks (\*) represent a statistical significance. DBH: diameter at breast height, SE: standard error

Despite careful observation, resprouting from the burnt bases of the *B. ermanii* stems was not observed in Plots 1 and 2 at 16 months post fire.

### Post-fire recovery of dwarf bamboos

The UAV-captured images revealed a rapid recovery of dwarf bamboos within 16 months post fire (Figures 5 and 6). Immediately after the fire, the aboveground portion was completely destroyed (Figure 1d). Six months post fire, the dwarf bamboos recovered to levels of 24.33% and 25.52% in Plots 1 and 2, respectively. Sixteen months post fire, the dwarf bamboo cover was 87.04% and 84.97% in Plots 1 and 2, respectively. The recovery rate at 10 months post fire was 62.71% and 59.45% in Plots 1 and 2, respectively (Figure 6). Field survey at 16 months post fire showed that the culm height was 67.8 ± 8.5 and 94.2 ± 7.0 cm in Plots 1 and 2, respectively.

**Figure 5.**
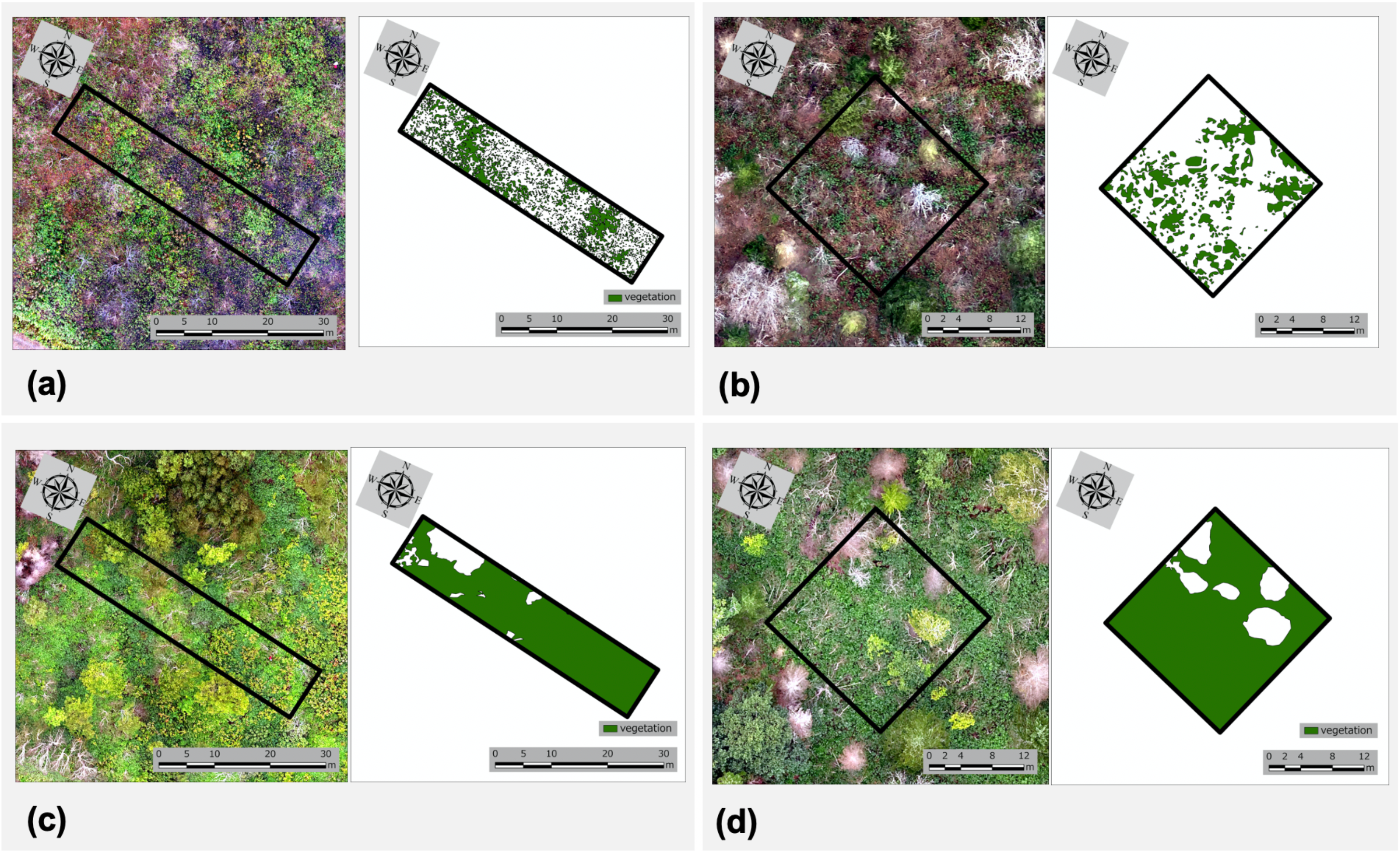
Map of *Sasa kurilensis* on the forest floor at 6 (2019) and 16 months (2020) after the fire in each plot. Mapping was manually interpreted from orthomosaic images captured with UAVs at each period.

**Figure 6.**
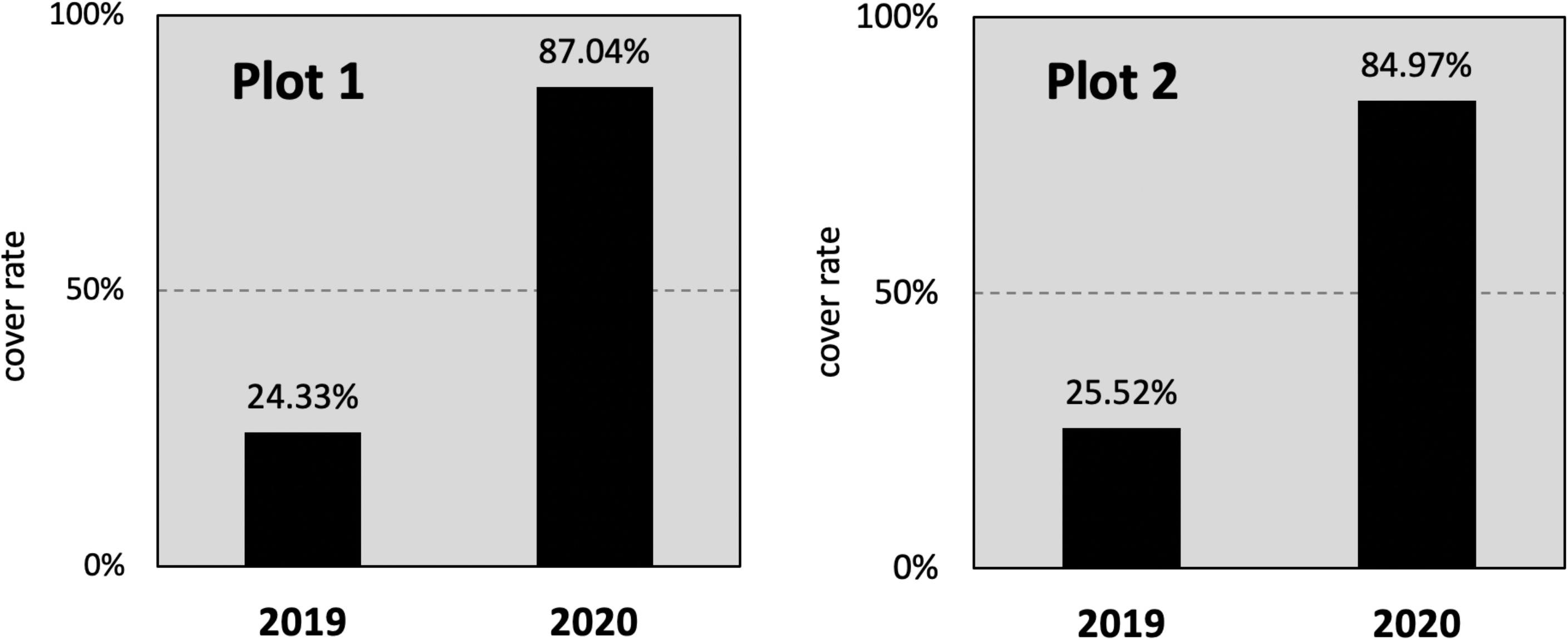
Cover rate of *Sasa kurilensis* estimated using UAV orthomosaic images.

### Post-fire vegetation and seedling emergence

Very few seedlings were established 16 months post fire in both plots (Table 4). The canopy tree species were *Abies sachalinensis* (Schmidt)*, Alnus japonica, Quercus crispula* Blume, *Phellodendron amurense* Rupr., *Cornus controversa* Hemsl., and *Salix bakko* Kimur*a.* In Plot 1, one individual each of *P. amurense*, *Acer pictum* Thumb. *mono* (Maxim.), and *Actinidia arguta* was recorded. In Plot 2, one individual each of *Aralia elata* and *A. arguta* was recorded. Seedlings of *B. ermanii* were not established in both plots.

**Table 4:** Summary of vegetation survey 16 months post fire.

|  | Stratum | Species | N |
| --- | --- | --- | --- |
| Plot 1 | Canopy | *Phellodendron amurense* Rupr | 3 |
|  |  | *Alnus japonica* | 2 |
|  |  | *Cornus controversa* Hemsl. | 1 |
|  |  | *Salix bakko* Kimura. | 1 |
|  | Subcanopy | *Sambucus racemosa* ssp. *kamtschatica* | 1 |
|  | Shrub | *Sasa kurilensis* | >100 |
|  |  | *Eupatorium chinensis* subsp. *sachalinens* | 12 |
|  |  | *Senecio cannabifolius* | 3 |
|  | Seedling | *Phellodendron amurense* | 1 |
|  |  | *Acer pictum* Thumb. *Mono* | 1 |
|  |  | *Actinidia arguta* | 1 |
| Plot 2 | Canopy | *Abies sachalinensis* (Schmidt) | 2 |
|  |  | *Quercus crispula* Blume | 2 |
|  | Subcanopy | - | - |
|  | Shrub | *Sasa kurilensis* | >100 |
|  | Seedling | *Actinidia arguta* | 1 |
|  |  | *Aralia elata* | 1 |

N: absolute shoot number

## Discussion

### Very low tree survival and non-resprouting of young forest stands

This study demonstrated the lethal effects of a fire on young *B. ermanii* trees within 2 years. The survival probability was clearly size-dependent in both plots, with larger trees (DBH > 8.9 cm) showing more tolerance to the fire in the first year (Table 2, Figures 3 and 4). Additionally, the relationship between fire tolerance and tree size has been recognised in fire-prone ecosystems such as the Siberian taiga (Uemura *et al.*, 1990) and Australian eucalyptus-dominated forests (Gill and Ashton, 1968). A previous study on *B. ermanii* trees in cool-temperate broad-leaved forests reported that many trees tended to die within a few years even if their canopies survived immediately after a forest fire (Sasa *et al.*, 1992). The results of these previous studies and the present study support the hypothesis that young forests are vulnerable to fire.

Post-disturbance resprouting is considered a common feature of *Betula* species (Perala and Alm, 1990; de Groot and Wein, 2004). In addition, based on a survey of old *B. ermanii* trees (probably >100 years) in the forest limit of high mountains in Central Japan, Okitsu (1991) reported that the percentage of resprouting (multiple-stemmed tree) *B. ermanii* trees was 25%–50%. This suggested that *B. ermanii* trees have the potential to resprout and play an important role in individual persistence. In contrast, in the present study, post-fire resprouting was not evident in young *B. ermanii* trees (<30 years) in both years. In particular, sprouting was not observed even from surviving individuals (DBH > 8 cm) at 6 months post fire, indicating that there was no response to fire regardless of survival. Thus, the low survival rate and lack of resprouting among the young trees demonstrate their vulnerability to fire damage and inability to allocate resources towards resprouting. Moreover, this phenomenon indicates that young *B. ermanii* trees are difficult to regenerate via resprouting post fire.

### Rapid recovery of dwarf bamboo and regeneration failure of young birch trees

We quantified dwarf bamboo coverage in both plots at 6 and 16 months post fire and found that the forest floor was densely and exclusively re-covered within 16 months post fire (Figures 5 and 6). Dwarf bamboos can dominate the forest floor through asexual reproduction after less severe disturbances (Sasa *et al.*, 1992; Matsuo et al. 2018). In the present study plots, the litter was only partially charred on the surface (Table 1). These results suggest that the fire caused relatively negligible damage to the belowground parts of the dwarf bamboos and showed strong resilience via vegetative reproduction in a short term than previously thought.

In addition to forest floor cover and recovery rates, the culm height of dwarf bamboos in both plots recovered by more than 50 cm (maximum height: 138 cm; Supplemental Figure 1) despite their complete destruction by the fire (Figure 1d). *Sasa kurilensis* cover is especially thick, where the culms reach a height of over 2 m (Oshima, 1962; Noguchi and Yoshida, 2005). In such areas, thick and dense dwarf bamboo covers block the light to seedlings and inhibit tree regeneration by preventing seedling establishment and suppressing seedling growth (Kobayashi *et al.*, 2004). Moreover, canopy removal by a fire is a critical mechanism that changes the structure and composition of forest vegetation (Mallik, 2003; Bond and Keeley, 2005; Dantas *et al.*, 2016). For example, in boreal and fire-prone temperate conifer forests, fire disturbances have been reported to cause regeneration failure for a long term due to rapid vegetative growth of understory ericaceous plants such as *Kalmia angustifolia* L (Mallik, 2003) and *Vaccinium myrtillus* L. (Mallik and Pellissier, 2000). These shrubs show a positive response to canopy gaps under higher light environments. Similarly, our UAV survey of dwarf bamboos suggests that the species can not only recover the aboveground parts immediately after the fire via the underground culm but also rapidly recover the aboveground biomass lost due to forest canopy removal (Figures 5 and 6).

Our findings of a high rate of mortality and lack of resprouting elucidated the vulnerability of young birch stands to fire. This indicates that young *B. ermanii* trees are unlikely to resprout for post-fire regeneration. Moreover, *B. ermanii* seedlings were not evident 16 months post fire under dense dwarf bamboo cover (Table 4). These results suggest that *B. ermanii* trees fail to successfully regenerate via seeds and resprouting under dense dwarf bamboo cover. In the dense dwarf bamboo understory, the emergence of tree species was almost inhibited regardless of the dispersal of new seeds (Nakashizuka, 1988; Doležal *et al.*, 2009). In addition, increased dwarf bamboo biomass reduces tree seedling density and plant species diversity in various forest understories (Nakashizuka, 1988; Hiura *et al.*, 1996; Narukawa and Yamamoto, 2002; Noguchi and Yoshida, 2004). Therefore, here, the dense and thick cover of dwarf bamboos that recovered post fire continued to inhibit the establishment of tree seedlings, resulting in regeneration failure.

### Implications for management

The vulnerability of young trees to fire demonstrates the importance of appropriate post-fire management in scarification sites even in cool-temperate broad-leaved forests. Dwarf bamboos in the study plots were largely removed by soil scarification in 1989. This species later invaded the soil scarification sites and recovered the forest floor with a resilience sufficient to tolerate the current level of fire disturbance for the next 30 years. Here, according to the UAV orthoimages and field survey (Supplemental Figure 1), many large and mature (mother) trees of *B. ermanii* (height >10 m and DBH > 40 cm; Masato Hayamizu’s personal observation) were observed around Plots 1 and 2. It is known that birch seeds are typically dispersed over relatively long distances; however, the closer the tree is to a mature tree, the greater the supply of birch seeds (Perala and Alm, 1990). Therefore, removing dwarf bamboos as soon as possible after a fire (e.g. early post-fire soil scarification) is an option for forest management through reforestation.

Soil scarification is a commonly used ‘close to nature’ forest management method that assists the natural regeneration of woody plants in various forest biomes, with widespread research globally, including Europe (Hynynen *et al.*, 2010; Nilsson *et al.*, 2010; Jäärats *et al.*, 2012), the United States (Woolley *et al.*, 2012), and Canada (Beaudry *et al.*, 1997; Giasson *et al.*, 2006), with recent reports from South–Central Chile (Soto and Puettmann, 2018), Lebanon (Nakhoul *et al.*, 2020), and Japan (Umeki, 2003). Globally, the concept of ‘close to nature’ has garnered attention, and it aims to reconcile wood production and ecological resilience (Messier *et al.*, 2013; O’Hara, 2016). Furthermore, soil scarification is of importance in this context. During recent years, several studies have focused on the spatial distribution (Ito *et al.*, 2019 a) and association among soil physicochemical properties (Ito *et al.*, 2019 b) of *B. ermanii* stands in soil scarification sites, because of an increasing commercial demand for Japanese birch trees (Ito *et al.*, 2018). However, information on the response and risk management to disturbances, including fire, in young forests post soil scarification is limited. The vulnerability of young forests and the factors involved in regeneration failure identified in this study suggest that future studies should focus on post-disturbance responses and risk assessment of young, dense stands in soil scarification sites.

## Conclusion

This study demonstrated the negative effects of a forest fire on young *B. ermanii* trees and elucidated the factors that contribute to the post-fire natural regeneration failure of young and dense stands in scarification sites. Young tree survival strongly depends on the tree diameter (DBH), and all birch trees died within 16 months of the fire. Moreover, (i) no resprouting was observed in any individual of *B. ermanii* trees, (ii) the rapid recovery of the dwarf bamboos covered nearly 90% of the forest floor within 16 months post fire, and (iii) a few seedlings of woody plant species emerged in forest understory covered by dense dwarf bamboos, suggesting that natural regeneration failure will continue in post-fire soil scarification sites. In particular, differences in fire tolerance and resilience between *B. ermanii* trees and dwarf bamboos are a major factor resulting in rapid vegetation change in cool-temperate broad-leaved forests in northern Japan. An evaluation of post-disturbance responses of young stands is necessary to understand whether this trend varies across regions and species.

## Funding

This work was supported by a research fund of the Hokkaido Research Organization.

## Acknowledgements

We thank the Hokkaido Government Okhotsk General Subprefectural Bureau Western Forestry Management for providing the data. We thank Ogura Takuro for assistance in the GIS analysis and Ebina Masuto for assistance and sharing valuable information. We thank Editage for English language editing.

## Conflicts of Interests

The authors declare that they have no known competing financial interests or personal relationships that could influence the work reported in this paper.

## Data Availability Statement

Data of this study will be available from the corresponding author upon request.

## Supplementary material

**Supplemental Table 1:**
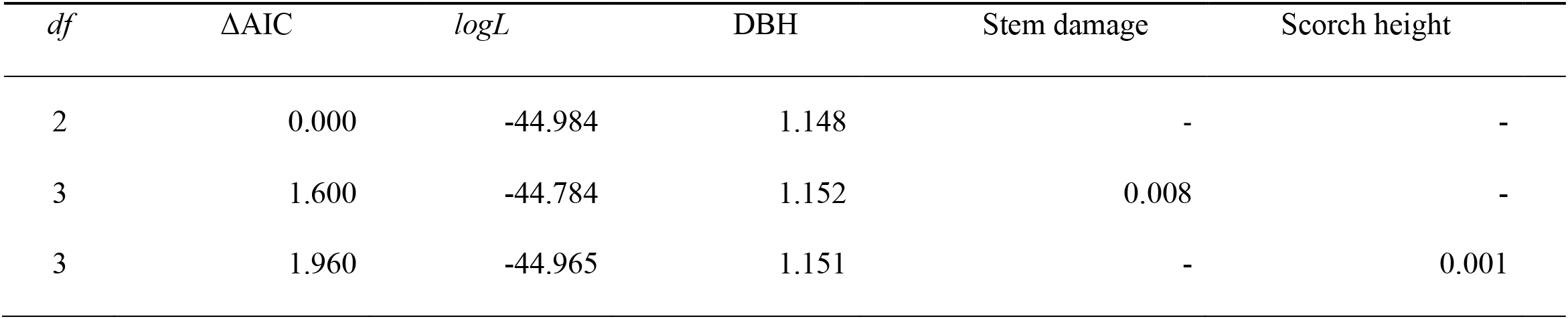
Results of GLM analysis on the effects of DBH, Stem damage, and Scorch height on probability of post fire survival of *Betula ermanii*. The terms with hyphen indicate that they were not included in the models by model selection procedure.

**Supplementa1 Figure 1:**
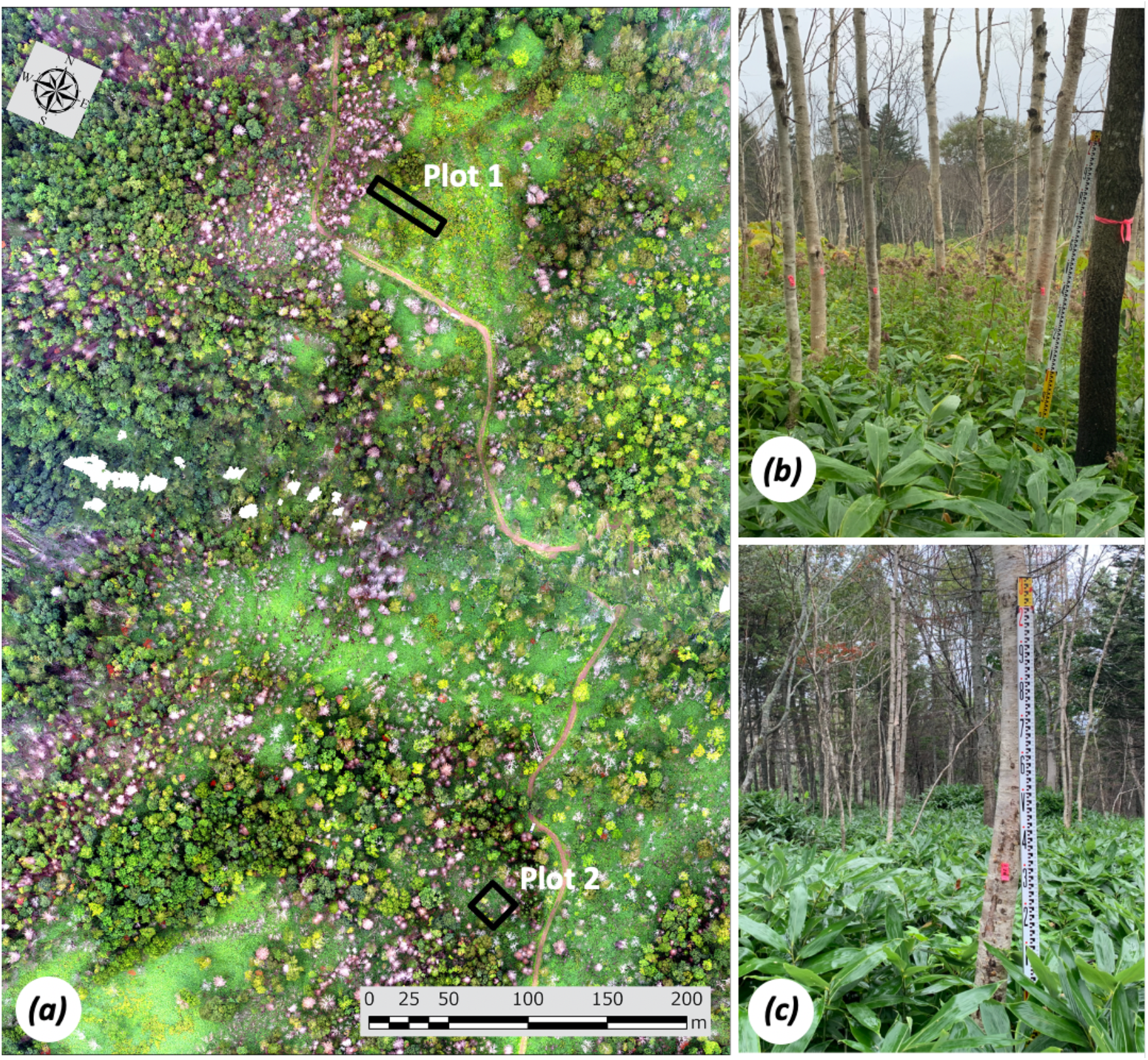
Orthomosaic images of *Betula ermanii* stands in scarification sites in Plots 1 and 2 in 2020, 16 months post fire, using UAV (a). *Betula ermanii* stands covered by dwarf bamboo in scarification sites after 6 months in Plot 1 (b) and Plot 2 (c). The maximum culm height of dwarf bamboos were 94 cm in Plot 1 and 138cm in Plot 2, as indicated by the Level staff in the photo (b and c).

## Notes

### Competing Interest Statement

The authors have declared no competing interest.

